# *SimBu*: Bias-aware simulation of bulk RNA-seq data with variable cell type composition

**DOI:** 10.1101/2022.05.06.490889

**Authors:** Alexander Dietrich, Gregor Sturm, Lorenzo Merotto, Federico Marini, Francesca Finotello, Markus List

**Author notes:** Joint last author (Francesca Finotello, Markus List).

## Abstract

**Motivation:** As complex tissues are typically composed of various cell types, deconvolution tools have been developed to computationally infer their cellular composition from bulk RNA sequencing (RNA-seq) data. To comprehensively assess deconvolution performance, gold-standard datasets are indispensable. Gold-standard, experimental techniques like flow cytometry or immunohistochemistry are resource-intensive and cannot be systematically applied to the numerous cell types and tissues profiled with high-throughput transcriptomics. The simulation of ‘pseudo-bulk’ data, generated by aggregating single-cell RNA-seq (scRNA-seq) expression profiles in pre-defined proportions, offers a scalable and cost-effective alternative. This makes it feasible to create *in silico* gold standards that allow fine-grained control of cell-type fractions not conceivable in an experimental setup. However, at present, no simulation software for generating pseudo-bulk RNA-seq data exists.

**Results:** We developed *SimBu*, an R package capable of simulating pseudo-bulk samples based on various simulation scenarios, designed to test specific features of deconvolution methods. A unique feature of *SimBu* is the modelling of cell-type-specific mRNA bias using experimentally-derived or data-driven scaling factors. Here, we show that *SimBu* can generate realistic pseudo-bulk data, recapitulating the biological and statistical features of real RNA-seq data. Finally, we illustrate the impact of mRNA bias on the evaluation of deconvolution tools and provide recommendations for the selection of suitable methods for estimating mRNA content.

**Conclusion:** SimBu is a user-friendly and flexible tool for simulating realistic pseudo-bulk RNA-seq datasets serving as *in silico* gold-standard for assessing cell-type deconvolution methods.

**Availability:** *SimBu* is freely available at https://github.com/omnideconv/SimBu as an R package under the GPL-3 license.

**Supplementary information:** Supplementary data are available at *Bioinformatics* online.

## 1 Introduction

The quantification of the cellular composition of complex tissues and organs is key to understanding pathophysiological changes underlying complex diseases like cancer, neurodegenerative diseases, and infections. For instance, the presence and abundance of specific immune cell types in the tumor microenvironment is associated with patients’ outcome and response to therapy (Fridman *et al*., 2012). Neurodegenerative diseases such as Alzheimer’s disease show evidence of cell pattern changes in the affected brain tissue (Johnson *et al*., 2020). Similarly, immuno-suppressive drugs such as corticosteroids can alter the composition of CD8+ T cells in cancer patients (Tokunaga *et al*., 2019).

Experimental methods such as fluorescence-activated cell sorting (FACS) or immunohistochemistry (IHC) can be used to estimate cell-type fractions within a sample using a predetermined set of markers (Petitprez *et al*., 2018), but they can only detect a limited number of cell types simultaneously. Also the estimation of cell-type proportions using single-cell technologies is not always feasible as it is too labor and cost intensive to be applied routinely or at a larger scale, and it can be affected by cell-type-specific bias in capture efficiency (Lambrechts *et al*., 2018). In contrast, widely available bulk RNA sequencing (RNA-seq) (Ozsolak *et al*., 2011) data from heterogeneous tissue samples can be analyzed *in silico* with deconvolution methods, which infer the cellular composition (i.e. cell-type fractions) of complex cell mixtures by leveraging cell-type-specific gene signatures (Sturm *et al*., 2019). These methods model the expression of a gene in one RNA-seq sample as the linear combination of its expression levels in the admixed cell types, weighted by their relative proportions (Finotello *et al*., 2018). Cell-type-specific gene signatures can either be derived from bulk RNA-seq data of sorted cells or from cell-type annotated single-cell RNA-seq data (scRNA-seq) (Travaglini *et al*., 2020; Hao *et al*., 2021).

A systematic benchmarking of deconvolution methods requires proper validation datasets. FACS and IHC can be used to generate gold-standard measurements of cell-type fractions from the same samples subjected to RNA-seq, but lack flexibility and scalability. In contrast, by simulating RNA-seq data artificially, gold-standard datasets with known cell-type fractions can be created and used to test deconvolution tools in multiple scenarios. As deconvolution tools are applied in diverse tissue and disease settings, their performance requires a frequent re-assessment. For benchmarking purposes, simulated ‘pseudo-bulk’ samples with controlled cellular compositions can be created by aggregating scRNA-seq profiles from cells having the same phenotype, either at the level of sequenced reads (Schelker *et al*., 2017; Finotello *et al*., 2019) or processed expression count data (Sturm *et al*., 2019; Cobos *et al*., 2020). To the best of our knowledge, a user-friendly and flexible tool for simulating pseudo-bulk data is currently unavailable, with the exception of a few basic functions available in the *immunedeconv* package (Sturm *et al*., 2019). Moreover, previous strategies neglect the potential bias of total mRNA amount, which can differ considerably by cell type (Islam *et al*., 2011) or state (Vallejos *et al*., 2017). Cells with lower (higher) mRNA content show lower (higher) total counts and number of expressed genes (Supplementary Fig. 1) and can be, thus, underestimated (overestimated) by deconvolution methods. Hence some deconvolution tools have incorporated scaling factors, which are used to correct the estimated cell-type fractions. *quanTIseq*, for instance, considers the median expression of the Proteasome Subunit Beta 2 (PSMB2) housekeeping gene (Finotello *et al*., 2019) across immune cell types. In contrast, *EPIC* (Racle *et al*., 2017) and *ABIS* (Monaco *et al*., 2017) use experimentally-derived scaling factors using FACS. However, these scaling factors are only available for a handful of immune cells from human tissues and are, thus, not applicable for general use.

Efforts have also been made to measure total mRNA levels in single cells, for instance using global reverse transcription followed by quantitative PCR (Jonasson *et al*., 2020). A different approach, which relies on an additional step during library preparation, is the inclusion of external RNA spike-in molecules, as suggested by the External RNA Control Consortium (ERCC) (Lichun *et al*., 2011). As these exogenous RNAs are added in equal concentrations to all cells, a higher proportion of reads mapped to spike-in transcripts in a cell indicates a lower endogenous mRNA content (Brennecke *et al*., 2013), and *vice versa*. Following this rationale, mRNA scaling factors consider the ratio of reads assigned to endogenous vs. exogenous (i.e. spike-in) molecules in each cell, (Vallejos *et al*., 2017), though they are not routinely available as they ‘consume’ a non-negligible portion of the sequencing depth, which could be instead fully dedicated to investigating the cell transcriptome. Due to this and additional limitations (Risso *et al*., 2014), other approaches derive scRNA-seq data-driven scaling factors by focusing on stably expressed genes over all cells as an alternative to spike-in RNAs (Lin *et al*., 2020) or estimate endogenous mRNA levels in single cells using a generative model based on the proportion between gene transcript counts and the relative frequencies of its mRNA, like the *Census* algorithm (Qiu *et al*., 2017). A systematic assessment of mRNA bias correction methods, however, is currently missing.

Here, we present *SimBu* (Simulating Bulk RNA-seq data), an R package that uses annotated scRNA-seq expression data to generate bias-aware pseudo-bulk RNA-seq datasets for benchmarking deconvolution tools (Fig. 1a). ScRNA-seq datasets can be obtained through the *Sfaira* (Fischer *et al*., 2021) database, which provides access to annotated scRNA-seq datasets from various organisms, tissues, and cell types. *SimBu* samples single cells from a scRNA-seq expression matrix and aggregates their transcriptomes to build pseudo-bulk RNA-seq expression profiles, summarized as gene counts, counts per million (CPM), or transcripts per millions (TPM), depending on the input data (Fig. 1b). SimBu is the first pseudo-bulk simulator that accounts for mRNA content bias; as cell types differ in mRNA contents (Supplementary Fig. 1), *SimBu* scales single-cell expression profiles using cell-or cell-type-specific mRNA factors. *SimBu* allows simulating pseudo-bulk datasets with diverse cell-type composition scenarios (Fig. 1c) to systematically assess specific features of deconvolution tools (e.g. accuracy, robustness to background noise, detection limits) (Sturm *et al*., 2019). In this work, we first demonstrate that *SimBu* can use Smart-seq2 or 10x Genomics scRNA-seq data to generate pseudo-bulk RNA-seq data that faithfully reflects the biological and statistical features of true bulk RNA-seq data. Second, we apply *SimBu* to systematically assess the performance of data-driven total mRNA scaling factors. *SimBu* code and documentation is available at https://github.com/omnideconv/SimBu under the GPL-3 license.

**Fig. 1.**
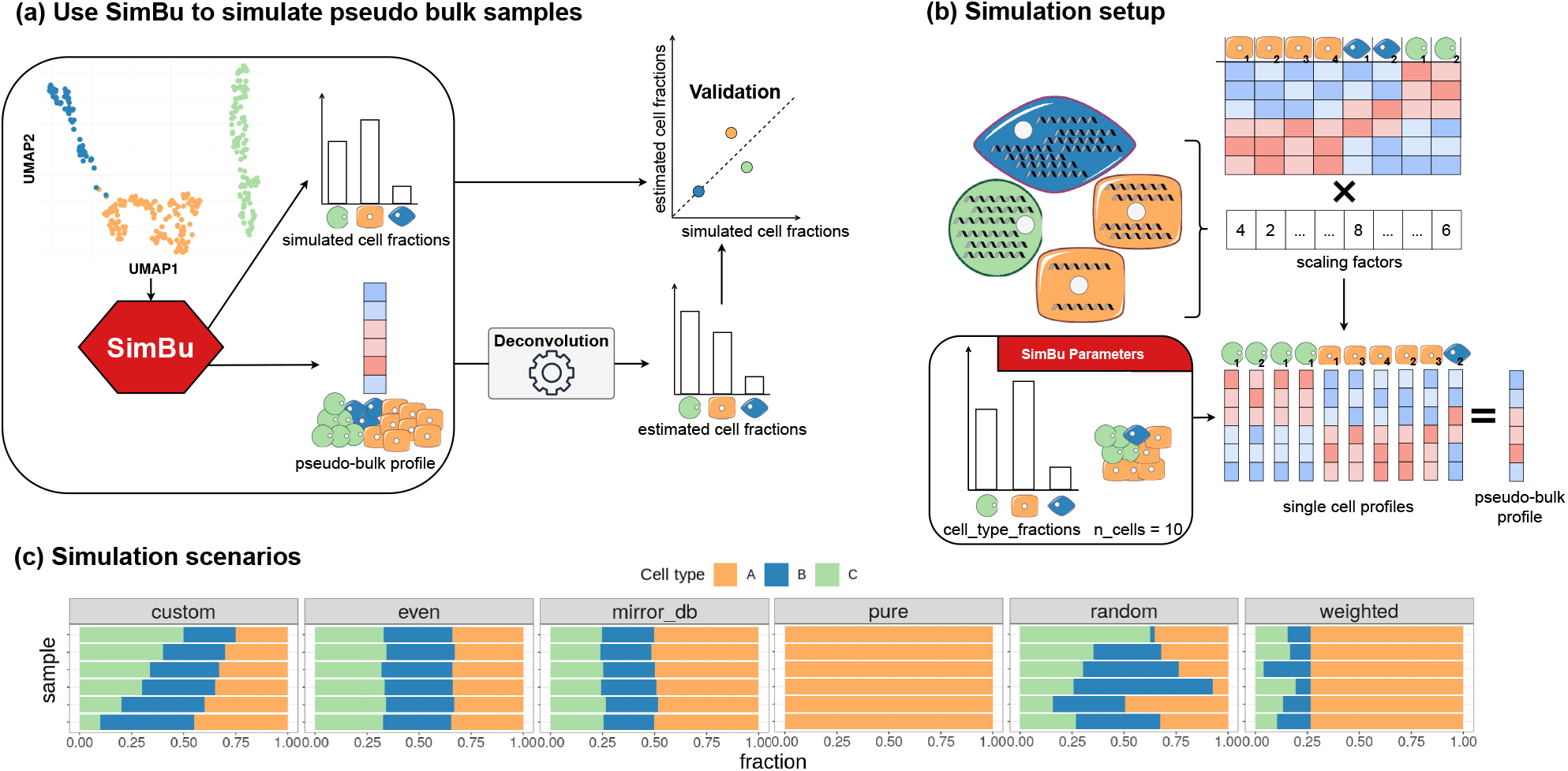
(a) SimBu uses cell-type annotated scRNA-seq transcriptomic data to generate RNA-seq expression profiles from pseudo-bulk samples with known cellular composition, which can be used to benchmark deconvolution tools. (b) SimBu takes as input a matrix of expression profiles from single cells and (optionally) multiplies them by different scaling factors to model cell-type-specific mRNA bias. Single-cell profiles are sampled randomly based on the specified cell-type fractions (`cell_type_fractions` argument) and the expected number of total cells in the sample (`n_cells`) and are aggregated into a single pseudo-bulk expression profile. (c) SimBu offers five different pre-defined simulation scenarios, with cell fractions specified by the user (custom), comparable in all cell types (even), mirroring the scRNA-seq dataset composition (mirror_db), representing a single cell type (pure), randomly distributed between all cell types (random), and consisting of one controlled cell type with controllable weight (weighted).

## 2 Methods

### 2.1 Input data and preprocessing

*SimBu* uses scRNA-seq expression data with matching cell-type annotation as mandatory input: raw count data is provided as a matrix of count values over *n* genes and *m* cells, together with a table describing cell-type annotations of all *m* cells present in the count matrix. *SimBu* can additionally take normalized Smart-seq2 or 10x scRNA-seq data as TPM or CPM, respectively, to account for differences in sequencing depth and/or gene length. For simplicity, normalized expression data are simply referred to as TPM throughout the manuscript. In addition, *SimBu* allows selecting annotated scRNA-seq data available through the *Sfaira* database (Fischer *et al*., 2021). Starting from the input scRNA-seq data, an object from the *SummarizedExperiment* (Morgan *et al*., 2021; Huber *et al*., 2015) package is built. Data can be loaded directly from h5ad files or *Seurat* (Hao *et al*., 2021) objects. The `filter_genes` and `variance_cutoff` filtering parameters can be used to remove genes with no expression or small variance over all cells respectively. Cell types can be removed if their abundance is below a specified threshold using the `type_abundance_cutoff` parameter. The blacklist and whitelist parameters give further control in selecting specific cell types.

### 2.2 Simulation based on custom or pre-defined scenarios

*SimBu* implements five pre-defined simulation scenarios, depending on the settings specified by the user through the `scenario` parameter (Fig. 1c):

- **custom**: cell-type fractions are defined by the user.
- **even**: all cell types are present in equal (or comparable) percentages; the `balance_even_mirror` parameter controls the intra-sample variance of cell fractions.
- **mirror_db**: mirror cell-type fractions as in the scRNA-seq dataset.
- **pure**: a single cell type (identified by the `pure_cell_type` parameter) makes up 100% of the pseudo-bulk sample.
- **random**: cell-type fractions are randomly sampled from the uniform distribution and scaled to sum up to 1.
- **weighted**: a cell type of interest (identified by the `weighted_cell_type` parameter) is assigned to a specific cell fraction (`weighted_amount`), whereas all other cell types are modelled according to the random scenario.

### 2.3 Pseudo-bulk simulation strategy

Depending on the chosen scenario, cell-type fractions are computed for each sample and multiplied by the number of cells contained in each sample (`n_cells`) to calculate the number of single-cell profiles to be sampled (with replacement) for each cell type. The selected single-cell profiles are optionally scaled to account for mRNA bias (see next section) and summed up to create a one-dimensional vector *v*_*bulk*_, representing the pseudo-bulk sample (Fig. 1b). When a precise sequencing depth is specified (`total_read_counts`), pseudo-bulk samples with higher or lower counts are either rarefied (Sanders, 1968) or multiplied by a factor, respectively, to equal the requested sequencing depth. If TPM data are provided, the same approach is carried out with normalized expression data from the same single cells. The final TPM pseudo-bulk profile is scaled to sum up to 10^6^.

### 2.4 Introducing a mRNA bias into simulations

The key feature of *SimBu* is the creation of pseudo-bulk RNA-seq samples that model mRNA bias, computed using scaling factors (Fig. 1b). Four data-driven approaches for introducing such a bias at the single-cell level are included in the package and are controlled by the `scaling_factor` parameter. The first two methods, *read_number* and *expressed_genes*, use the number of mapped reads and the number of expressed genes per cell, respectively. Next, *Census* (Qiu *et al*., 2017), which is an algorithm implemented in the *Monocle2* package, estimates lysate transcript counts. We note that *SimBu* uses its own implementation of *Census*, to improve run-time efficiency but the algorithm itself is unchanged. Finally, *spike_in* considers external spike-in counts if available in the input scRNA-seq data. For each cell, this factor is computed as the ratio between the number of reads mapped to genes over total reads (i.e. the sum of gene and spike-in reads), therefore giving the ratio of reads that are not mapped to spike-ins. When a scaling factor is computed for individual cell types, the median scaling factor per cell type is applied (Supplementary Fig. 1). *SimBu* additionally offers the option to use pre-calculated scaling factors, i.e. *EPIC* (Racle *et al*., 2017), *quanTIseq* (Finotello *et al*., 2019) and *ABIS* (Monaco *et al*., 2017), which are already available as a single estimate per cell type. Scaling factors are multiplied with the expression matrix, to add bias to the cells. It is also possible to supply a custom scaling factor with the `custom_scaling_vector` parameter, where a list with scaling values of length *m* needs to be provided.

### 2.5 Bulk and single-cell RNA-seq data

We considered scRNA-seq data generated from mouse spleen by the *Tabula Muris* (The Tabula Muris Consortium, 2018) project, encompassing five cell types (B cells, T cells, NK cells, dendritic cells and macrophages) using two different single-cell platforms: Smart-seq2 (Picelli *et al*., 2014) and 10x Chromium (Zheng *et al*., 2017). We integrated the two datasets using the SCTransform workflow from the R package Seurat (Hao *et al*., 2021) and re-annotated T cells, selecting only the CD4, CD8 and T regulatory cells subtypes. In parallel, two publicly available bulk RNA-seq datasets containing mouse spleen data were used, called *Petitprez* (Petitprez *et al*., 2018) and *Chen* (Chen *et al*., 2017, 2018). The bulk RNA-seq datasets were accessed in March 2022 with the codes E-MTAB-9271 (*Petitprez*) and E-MTAB-6458 (*Chen*).

Additionally, we used three public human single-cell datasets as the basis for several simulations: the *Travaglini* dataset is a single-cell atlas for the human lung, where cells were processed using the 10x and Smart-seq2 assays (Travaglini *et al*., 2020). For this study, we only used the Smart-seq2 analyzed cells due to their deeper profiling. The resulting subset contains 8,657 cells with FACS annotated cell types. As only read counts are available, we calculated TPM counts using the available gene length information. Notably, this dataset contains ERCC spike-in counts for all cells. It was latest accessed in March 2022 and is available at https://www.synapse.org/#!Synapse:syn21041850.

Another Smart-seq2 dataset, the *Maynard* dataset, provides 20,592 cells from human metastatic lung (Maynard *et al*., 2020). Read counts and TPM counts are available, processed with the nf-core RNA-seq pipeline v3.0 (Ewels *et al*., 2020). We performed unsupervised clustering and manual cell-type annotation. This dataset was accessed in March 2022 with the accession code PRJNA591860.

The third dataset, *Hao*, contains human PBMC cells (Hao *et al*., 2021), analyzed with CITE-seq (Stoeckius *et al*., 2017), which produces comparable results to 10x assays. 161,764 cells are available, which are labelled using unsupervised clustering. As no gene length is available for 3’ end sequencing methods, we calculated CPM values instead of TPMs here. This dataset was accessed last in March 2022 with the accession code GSE164378.

### 2.6 Simulations

Based on the *Tabula Muris* dataset, we performed a set of simulations, which used the cell-type fractions and sequencing depth of the spleen samples in the *Petitprez* and *Chen* datasets. This resulted in three mouse pseudo-bulk samples based on the *Chen* dataset and four pseudo-bulk samples based on the *Petitprez* dataset. We used the cell-type fractions of the true bulk samples as the basis for these simulations. Only cell types which are present in both single-cell and bulk dataset could be used for these simulations (see Supplementary Table 3). The simulations were performed with and without an added mRNA bias, based on the number of expressed genes (Fig. 2, Supplementary Fig. 2-4). Each simulated sample contains 500 cells.

**Fig. 2.**
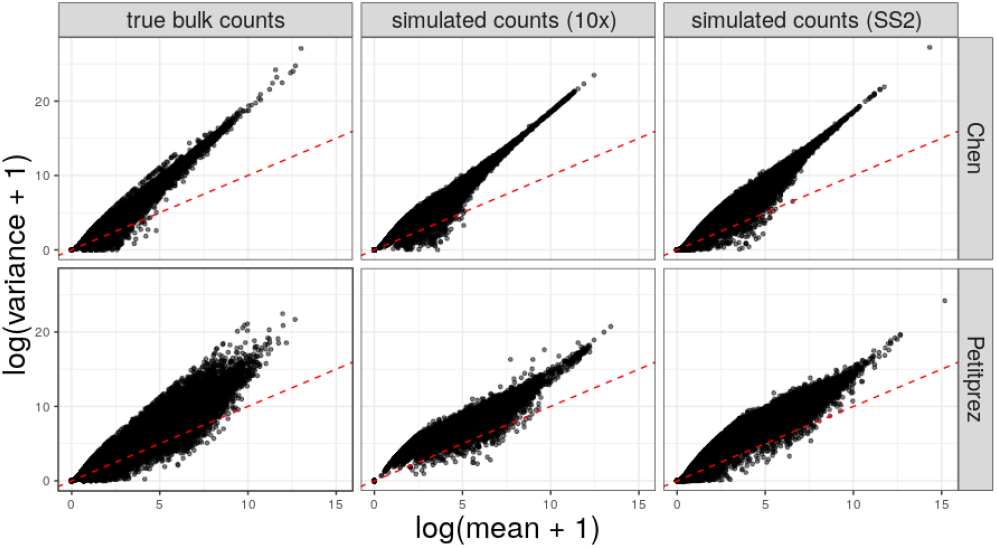
Comparison of gene-wise mean and variance of count data in different bulk and pseudo-bulk samples. The first column contains data of two true bulk datasets, Petitprez (Picelli et al., 2014) and Chen (Chen et al., 2017). Column two and three contain simulated pseudo-bulk data based on mouse spleen scRNA-seq data (Tabula Muris), generated with the 10x Chromium (10x) and Smart-seq2 (SS2) assay, respectively. Pseudo-bulk samples were generated using only those cell types present in both the true bulk sample and the single-cell dataset, mimicking the sequencing depth of the true bulk samples.

We performed a second set of simulations, using the three human scRNA-seq datasets, applying different calculations for the mRNA bias. In the first round, one out of four cell types (B cells, monocytes, natural killer cells, CD8+ T cells) was selected and an extreme bias of ten was applied to only them, while the remaining cells went untouched. Next, the previously described real world biases were added to all simulations, using the `scaling_factor` parameter in *SimBu*. The *Travaglini* dataset was the only one with spike-in counts, so we could calculate the `spike_in` scaling factor only for this dataset. In the described analyses, we focused on the immune cells in these datasets; *SimBu*’s `whitelist` parameter was used to keep these cells (Supplementary Table 1 and 2 for details on cell types). For both rounds of simulations, 100 pseudo-bulk samples were created with the random scenario, each containing 1,000 cells.

## 3 Results

### 3.1 *SimBu* generates valid pseudo-bulk data

RNA-seq read counts follow the negative binomial (NB) distribution (Robinson *et al*., 2010) and *SimBu* simulates counts faithfully, following the expected NB distribution with variance larger than the mean. We show this by comparing pseudo-bulk datasets generated from Smart-seq2 and 10x scRNA-seq data from the *Tabula Muris* project with two real bulk RNA-seq datasets (Fig. 2). Despite the fact that true bulk and pseudo-bulk do not contain exactly the same cell types (see Methods and Supplementary Table 3), dataset-specific characteristics, such as a less dispersed distribution of genes with high mean and variance in the *Chen* (Chen *et al*., 2017) dataset, are also recapitulated by the simulations. We could also confirm a strong, significant correlation between the pseudo-bulk data generated from 10x and Smart-seq2 scRNA-seq data at the level of gene counts and TPM (Spearman’s *ρ* = 0.9) (Supplementary Fig. 3 and 4). Compared to Smart-seq2 pseudo-bulk, 10x-based pseudo-bulk show a overestimation of highly-expressed genes and under-estimation of lowly-expressed genes (Supplementary Fig. 2), in agreement with the higher throughput of this technology (Baran-Gale *et al*., 2018). As *SimBu* can also apply mRNA scaling factors, which alter the single-cell expression profiles, we performed an additional set of simulations including a mRNA bias based on the number of expressed genes. Only a minor difference in the count behavior compared to those simulations without a bias could be noted, showing that our simulations are robust to possible statistical changes due to a scaling factor (Supplementary Fig. 2). Similarly, the addition of mRNA bias shows no impact on the linear relationship of count or TPM data between different assays (Supplementary Fig. 3 and 4).

### 3.2 Comparing different scaling factor approaches

We compared different approaches for estimating mRNA scaling factors in human immune cell types: two based on an experimental assay, one pre-calculated data-driven approach, and four data-driven approaches applied to three single-cell datasets (Fig. 3). Data-driven approaches can be used to estimate mRNA content at a single-cell level, whereas the values originating from experimental assays only give one estimate per cell type. For each pair of approaches, we considered only matching cell types, and used Spearman’s correlation to quantifying their agreement as we did not expect linear relationships between scaling factors.

**Fig. 3.**
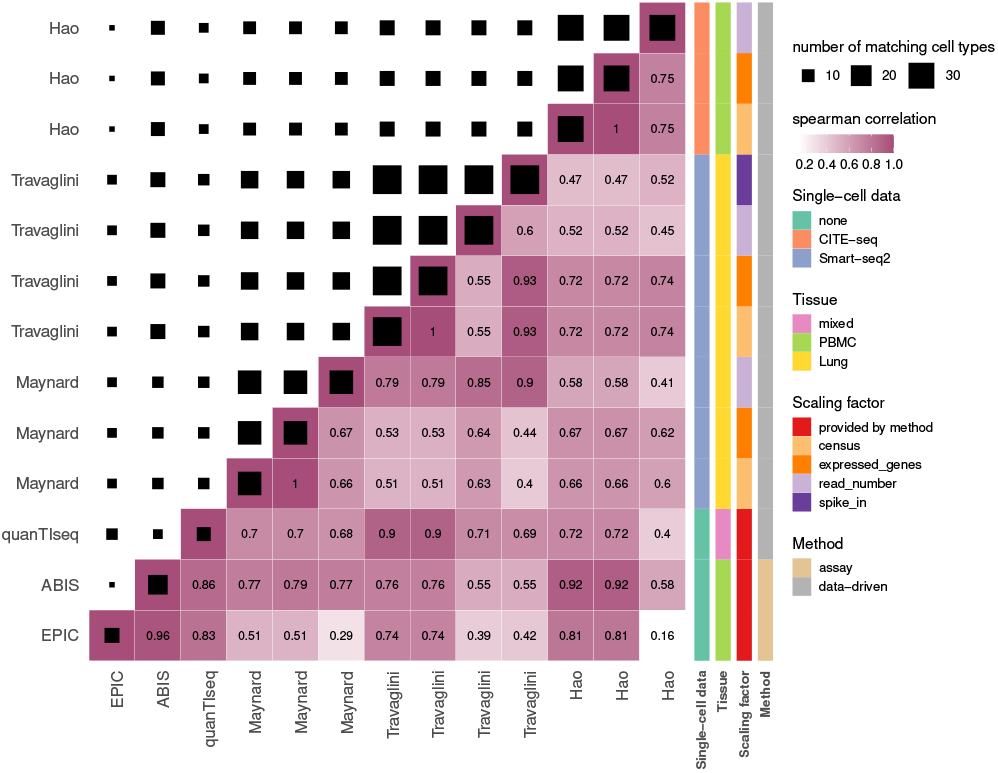
Comparison of different mRNA scaling factors. Upper triangular matrix shows the number of matching cell types for each pairing. Lower triangular matrix shows a Spearman correlation heatmap, including the Spearman correlation coefficient. Additional group indications are given by coloured boxes on the left: Method indicates if the scaling factor is derived by an experimental assay or data-driven. Scaling factor indicates the used calculation for the data-driven approaches or the name which uses an assay. Tissue gives the name of the tissue that was used in the respective approach. Single-cell data gives the name of the used single-cell assay of the corresponding dataset.

*EPIC* and *ABIS* scaling factors, which are based on experimental assays and showed a high correlation (*ρ*=0.96), hence we are considering both as gold-standard. Among the data-driven scaling factors, the number of expressed genes showed high correlation with experimental scaling factors in both the *Hao* (*EPIC*: Spearman’s *ρ*=0.81, *ABIS*: *ρ*=0.92) and *Travaglini* (*EPIC*: *ρ*=0.74, *ABIS*: *ρ*=0.76) dataset. The pre-calculated scaling factor derived by *quanTIseq* comes close to these methods (*EPIC*: *ρ*=0.83, *ABIS*: *ρ*=0.86), while expressing the highest correlation with the gene based scaling in the Travaglini dataset (*ρ*=0.9).

We observed a surprisingly high correlation between the number of expressed genes and *Census* factors in all three datasets on a cell type (Fig. 3) and single-cell level (Supplementary Fig. 5)(*ρ*=1). *Census* estimates endogenous mRNA levels in cells using TPM counts, which appears to result in a value that is almost identical to the number of expressed genes in a cell. *Travaglini* was the only available single-cell dataset with a wide range of well-annotated immune cells and spike-in count information (ERCC spike-ins). As explained in the introduction, spike-in derived scaling factors can be considered as a silver-standard for the content of mRNA in a cell (see Methods 2.4 for details on scaling factor calculation). When comparing this scaling factor on a single-cell level with the number of genes or reads as scaling factors, the former showed a noticeably higher correlation coefficient (*ρ*=0.876), indicating closer agreement between spike-in and gene-based scaling factors (Supplementary Fig. 5). In summary, mRNA scaling factors based on the number of expressed genes are strongly correlated with both, gold- and silver-standard scaling factor methods based on experimental evidence.

### 3.3 Use case: impact of mRNA bias on deconvolution

The primary goal of *SimBu* is to generate realistic pseudo-bulk RNA-seq profiles which can be used for benchmarking deconvolution tools. To showcase the value of generating mRNA bias-aware pseudo-bulk data, we performed two sets of simulations based on the *Travaglini* dataset with different scaling factors and analyzed them with different deconvolution methods to study the impact of mRNA bias. In the first scenario, we applied a custom, extreme scaling factor to a single cell type (see Methods 2.6 for details). Some noticeable differences could already be spotted in its influence on cell-type estimates (Fig. 4a and Supplementary Figure 6), where a higher overall mRNA content for a single cell type resulted in a systematic over-estimation of the corresponding cell-type fractions via deconvolution. Without moving into a methodical evaluation of deconvolution methods, we simply note that mRNA can manifest as a more subtle ‘spillover effect’ affecting closely related cell types. For instance, *quanTIseq* assigns monocyte signal to dendritic cells and macrophages (Fig. 4a), whereas *EPIC* and *CIBERSORT* overestimate natural killer cells (NK cells) in case of extreme CD8+ T cells mRNA bias (Supplementary Fig. 6, 9). In the PBMC based simulations (Supplementary Fig. 7), spillover effects were less prominent; only an increased signal of monocytes results in minor over-estimation of CD4+ T cells by *EPIC* and dendritic cells by *quanTIseq*. B cells appear to be more easily distinguishable by all deconvolution tools in all simulations, with only *EPIC* and *CIBERSORT* showing a spillover effect (mainly to T cell subtypes). This shows how B cell estimates can be affected to a higher degree by mRNA bias.

**Fig. 4.**
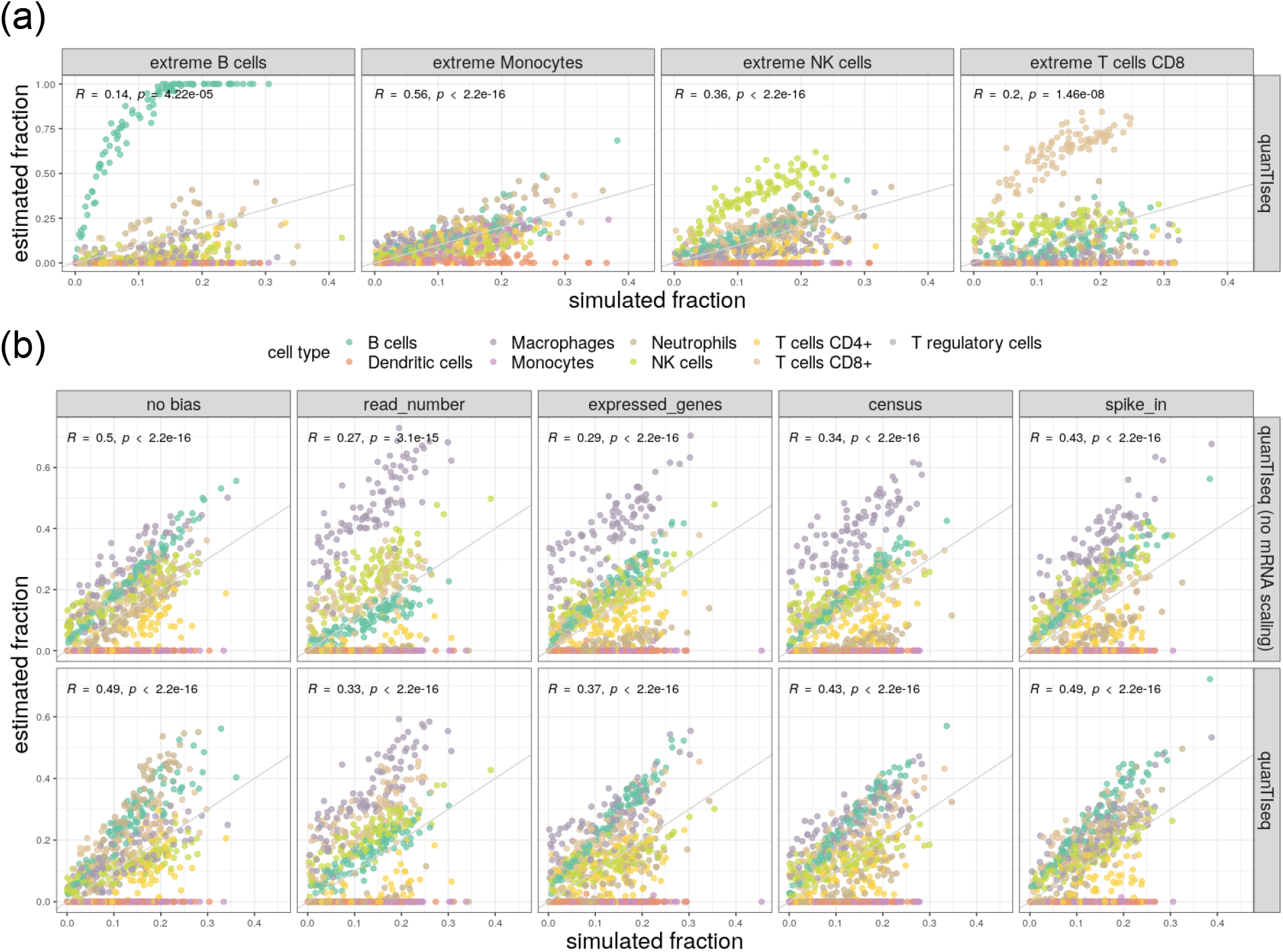
Effect mRNA bias on deconvolution of Travaglini based pseudo-bulk simulations. (a) An extreme scaling factor was applied to four selected cell types and quanTIseq was used to estimate cell-type fractions (y-axis). The x-axis shows the true, simulated cell-type fractions. (b) Comparison of estimated and simulated cell-type fractions with/without mRNA bias (computed with four different scaling factor methods) and with/without quanTIseq mRNA bias correction.

Starting from the same human scRNA-seq data, we assembled a second simulation to show the impact of modelling true biological mRNA bias. Pseudo-bulk data, generated with and without mRNA bias, was analyzed with *quanTIseq*, with and without its internal mRNA bias correction. In the absence of mRNA bias, this ‘uncorrected’ *quanTIseq* deconvolution resulted in an accurate estimation of most cell types, as the fractions of simulated and estimated cell types converge at the identity line (Fig. 4b). When instead ‘uncorrected’ *quanTIseq* was applied to pseudo-bulk datasets containing mRNA bias, this resulted in a systematic over-estimation of macrophages and under-estimation of neutrophils (Fig. 4b), as expected due to their marked mRNA content differences (Supplementary Fig. 1). The largest deviation from the identity line appears with a bias based on the number of mapped reads per cell (read_number), something that could be expected given its lower correlation to other methods (Fig. 3). By default, *quanTIseq* applies an internal mRNA bias correction with the goal of accounting for differences in mRNA levels in cell types. Performing the deconvolution with this setting helps to ameliorate systematic biases including the over-estimation of macrophages and under-estimation of neutrophils (Fig. 4b).

We could observe similar patterns in the *Hao* and *Maynard* datasets (Supplementary Fig. 8, 10). Additionally, in the *Maynard* simulations with real-world mRNA bias, macrophages were always over-estimated; only corrected deconvolution on simulations without bias could result in correct estimation of this cell type. The *Hao* simulations show how cell types such as monocytes and dendritic cells benefit from adding mRNA bias, with a less prominent over-estimation of both cell types. We could show in this scenario, how modelling pseudo-bulk RNA-seq samples using mRNA bias is of importance when testing deconvolution results.

## 4 Discussion

The computational deconvolution of the cellular composition of complex tissues from bulk RNA-seq has opened new avenues in the investigation of human diseases. The rapid advance of single-cell technologies has further expanded the scope of these computational methodologies to additional cell types, tissues, and organisms. However, the successful application of deconvolution relies on the possibility to comprehensively and systematically benchmark these methods on proper validation datasets. As it is not feasible to generate gold-standard data for every scenario of interest, we developed *SimBu*, the first comprehensive simulator of pseudo-bulk RNA-seq data, that uses scRNA-seq data to flexibly model various cell-type compositions in an mRNA bias-aware fashion.

Unlike previous approaches that did not account for cell-type-specific differences in mRNA content (Schelker *et al*., 2017; Sturm *et al*., 2019), *SimBu* models mRNA bias by introducing a scaling factor in the pseudo-bulk simulations, altering the single-cell expression profiles depending on the estimated amount of endogenous mRNA per cell type. *SimBu* allows the selection of various data-driven scaling factors.We tested these scaling factors using three human scRNA-seq datasets (Travaglini *et al*., 2020; Maynard *et al*., 2020; Hao *et al*., 2021) from different human tissues (lung and PBMC) and single-cell technologies (Smart-seq2 and 10x). Despite the caveats in the comparison of different cell types and tissues, most of scaling factors showed a good agreement with experimentally derived mRNA abundances and spike-in based scaling factors. A distinctive advantage of single-cell data-driven scaling factors is the possibility of deriving them from any scRNA-seq dataset, cell type, tissue and even organism. These approaches can be further extended to the quantification of mRNA abundance in different cell states (e.g., in resting vs. activated T cells or cells at different stages of their cell cycle). We demonstrated that *SimBu* can robustly generate pseudo-bulk simulations from 10x and Smart-seq2 scRNA-seq data with NB-distributed gene counts and highly correlated per-sample expression profiles at count and TPM level. By returning as output both counts and TPM pseudo-bulk profiles, *SimBu* allows evaluating deconvolution tools with different input-data requirements. We further demonstrated the impact of modelling mRNA bias by analyzing bias-included pseudo-bulk data with different ‘absolute’ deconvolution methods, which return cell fractions that are comparable between cell types (Sturm *et al*., 2019). The ‘extreme-bias’ scenario was helpful to illustrate that cell-type-specific mRNA can have a big, direct impact on the quantification of the corresponding cell types, but can also manifest in more subtle ways due to spillover effects between closely related cells types. By using *quanTIseq* as an example, we demonstrated that mRNA-bias aware simulations are necessary to ensure a fair evaluation of the ability of deconvolution methods to account for mRNA bias. When planning SimBu simulations of pseudo-bulk datasets, we recommend selecting the gold-standard mRNA scaling factors derived by *EPIC* or *ABIS* if the (immune) cell types of interest, are covered. While quanTIseq lacks experimental evidence for its scaling factors, it offers comparable results. As a second choice, in case other cell types needs to be considered, we recommend using spike-in based scaling factors, if these are available in the scRNA-seq data, to be used in the simulations. If neither of those approaches are applicable, single-cell data-driven scaling factors based on the number of expressed genes per cell portrays the closest representation of the true mRNA levels in cells. One advantage of single-cell data-driven scaling factors is the possibility to derive them from any scRNA-seq dataset, as we showed with the *Tabula Muris* based simulations, thus allowing the simulation of any cell types, tissues, and organisms of interest.

In this work, we illustrated the value of *SimBu* on “first-generation” deconvolution methods based on pre-computed signatures. Nevertheless, we envision that SimBu will be vital for benchmarking “second-generation” methods that are trained using scRNA-seq data (Newman *et al*., 2019). Thanks to its features, *SimBu* is well-suited to systematically and flexibly assess the applicability of these methods to different contexts. Nevertheless, purely scRNA-seq data-driven approaches show a good agreement with those including experimental evidence and therefore can be used from any single-cell dataset, as we showed with the *Tabula Muris* based simulations. Beyond RNA-seq, we plan to adapt the *SimBu* simulation framework to other assays, e.g. DNA methylation, spatial transcriptomics or Assay of Transposase Accessible Chromatin sequencing (ATAC-seq), were gold standard pseudo-bulk datasets are also needed to study cell type deconvolution.

## Supporting information

scRNAseq_celltypes

## 5 Availability

*SimBu* is available via GitHub at https://github.com/omnideconv/SimBu as an R package under the GPL-3 license and comes with detailed documentation vignettes to aid in the setup of pseudo-bulk simulations.

## Acknowledgements

We would like to thank Katharina Reinisch and Constantin Zackl for testing *SimBu* and providing feedback on the implementation and documentation.

## Funding

This work was supported by the German Federal Ministry of Education and Research (BMBF) [031L0294A to M.L.], by the Austrian Science Fund (FWF) [T 974-B30 to F.F.] and by the Oesterreichische Nationalbank (OeNB) [18496 to F.F.]. GS was supported by a DOC-fellowship from the Austrian Academy of Sciences.

## Notes

### Competing Interest Statement

The authors have declared no competing interest.

