## Supplementary Figures and Tables for "*SimBu*: Bias-aware simulation of bulk RNA-seq data with variable cell type composition"

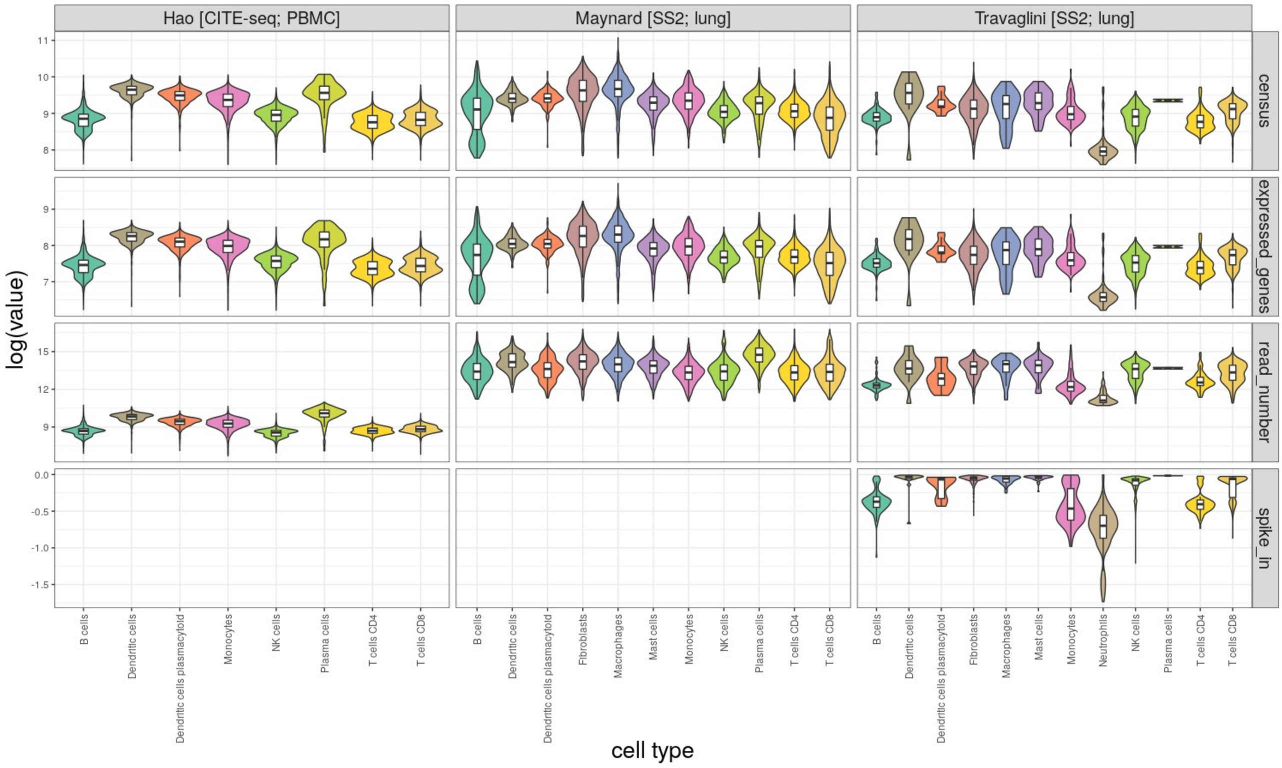


**Supplementary Figure 1:** Violin plot for mRNA content estimations using four different data-driven methods on three scRNA-seq datasets. Note that only the *Travaglini*[3] dataset had spike-in counts available, so its the only one where the spike_in scaling factor could be calculated. Also, this only shows a selection of cell types, as more cell types were annotated for each dataset.

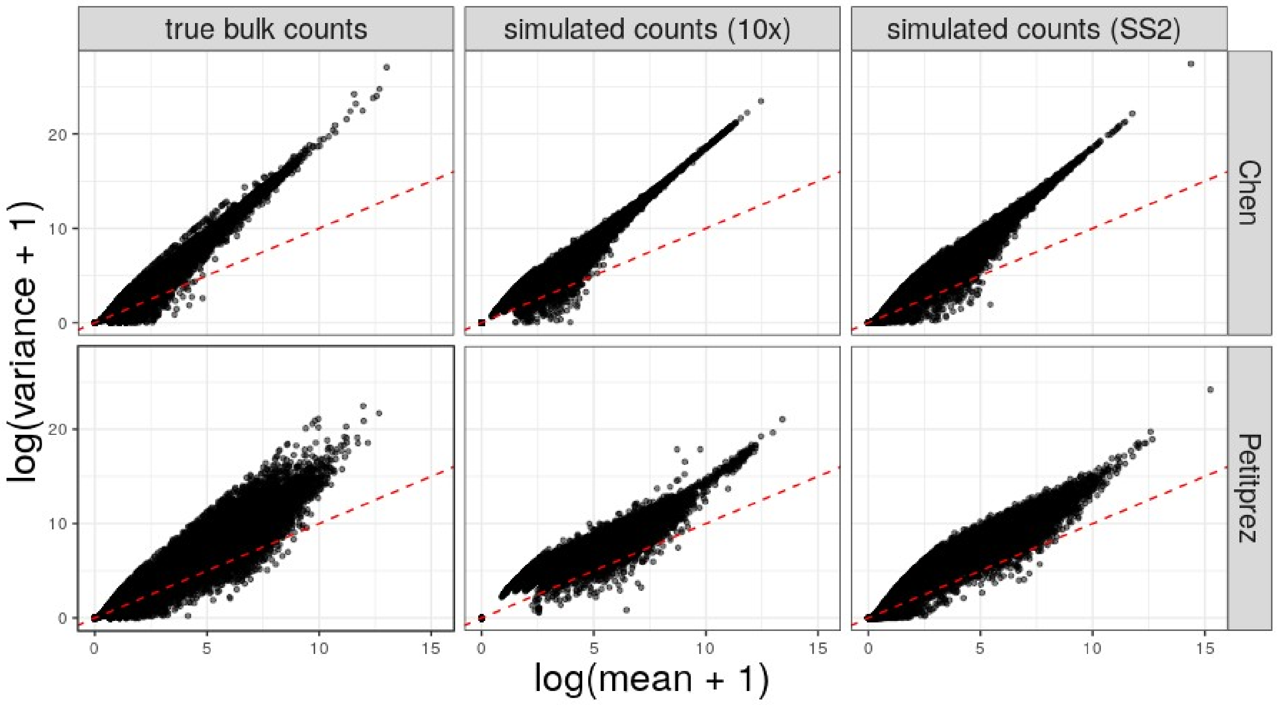


**Supplementary Figure 2:** Comparison of gene-wise mean and variance of count data in different bulk and pseudo-bulk samples. Same setup as in Figure 2 of main text, but with additionally added mRNA bias based on the number of expressed genes per cell.

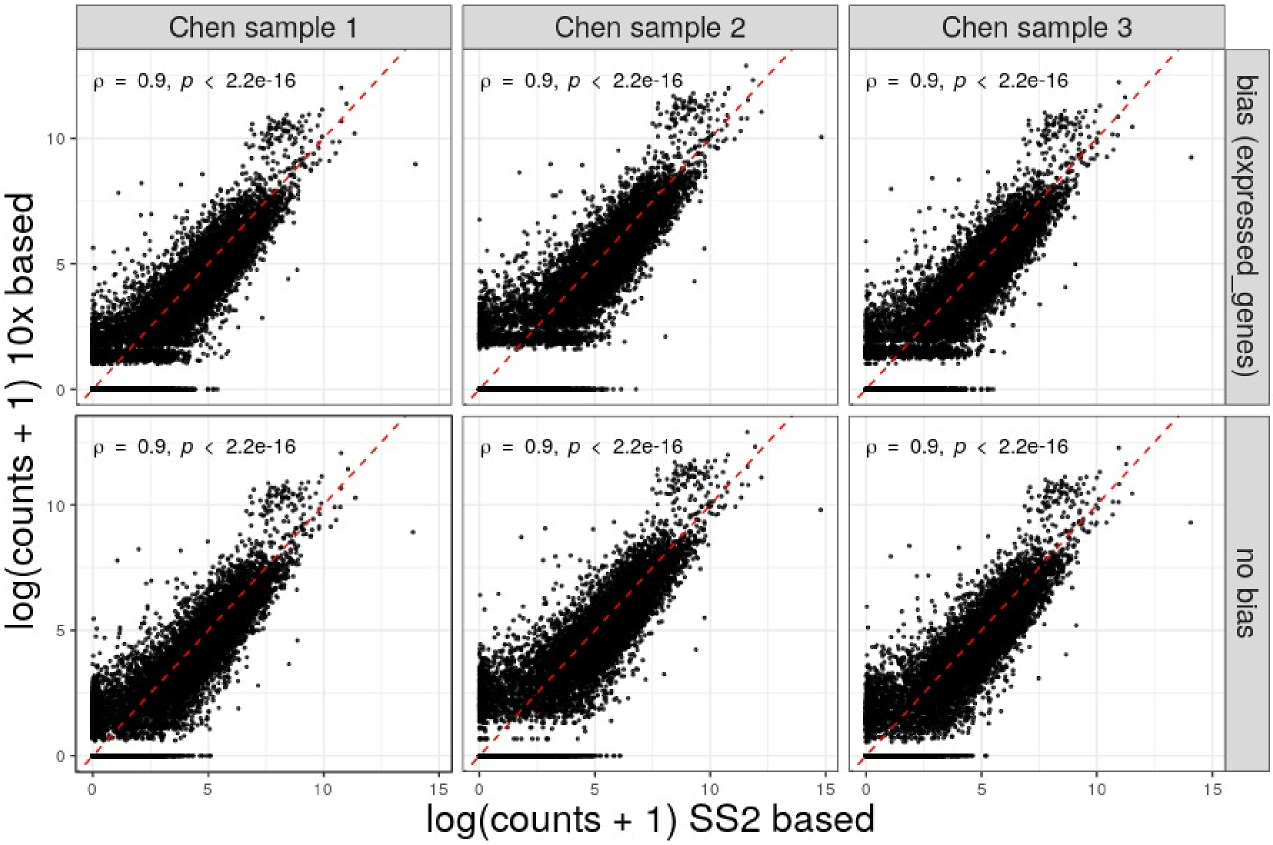


**Supplementary Figure 3:** Comparison of simulated counts based on the Smart-seq2 and 10x Chromium based scRNA-seq datasets of Tabula Muris. Upper row shows simulated counts with additionally added mRNA bias based on the number of expressed genes per cell.

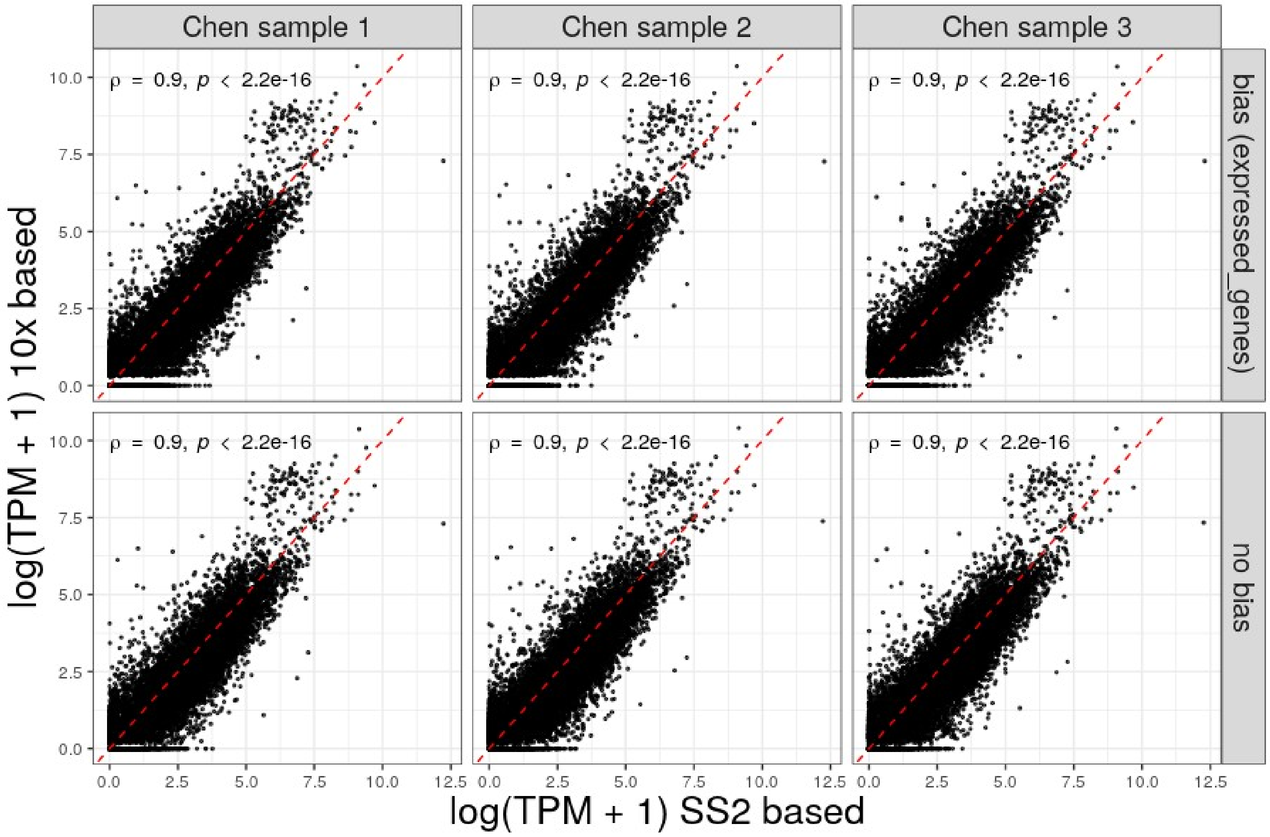


**Supplementary Figure 4:** Comparison of simulated TPMs based on the Smart-seq2 and 10x Chromium based scRNA-seq datasets of Tabula Muris. Upper row shows simulated counts with additionally added mRNA bias based on the number of expressed genes per cell.

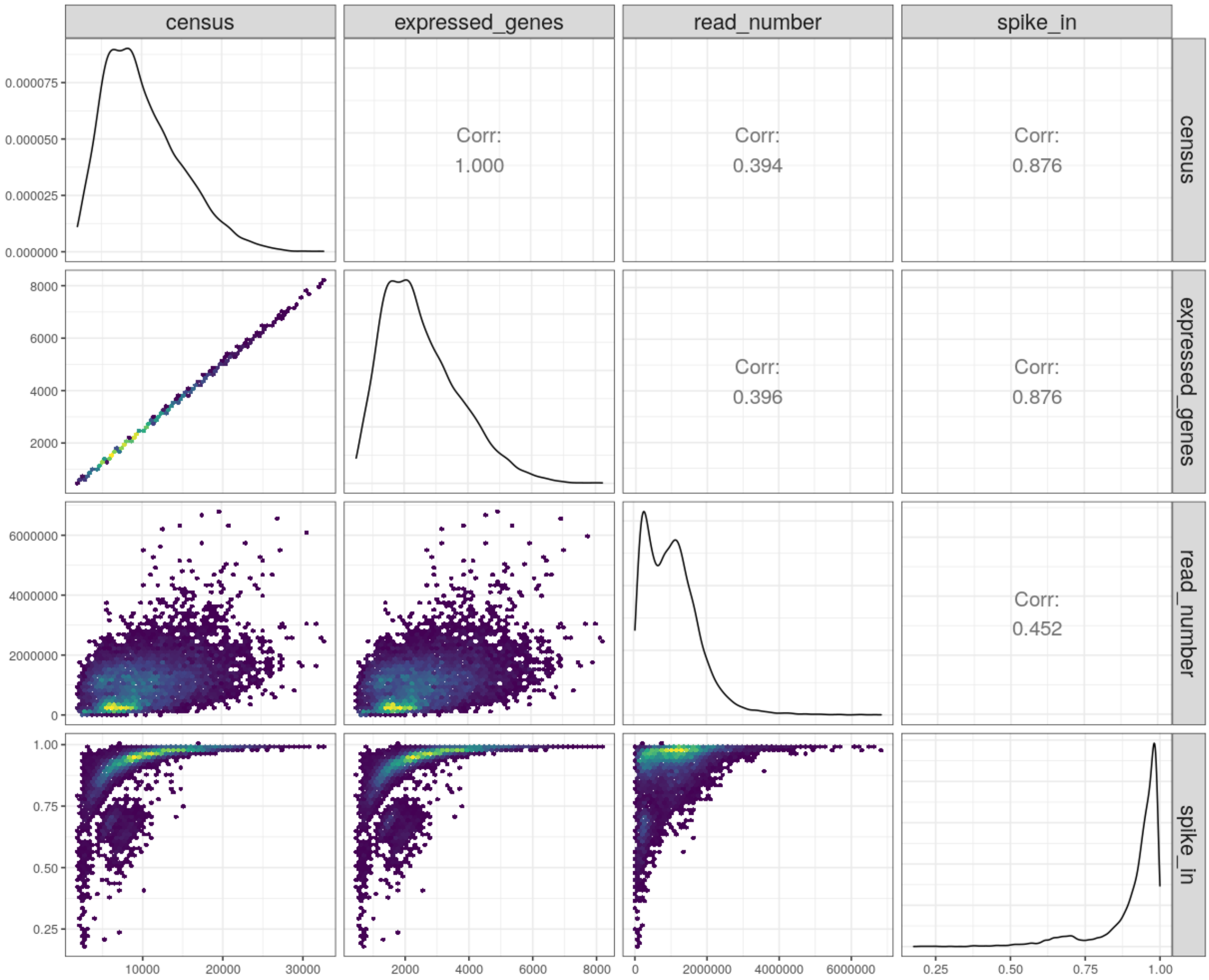


**Supplementary Figure 5:** Distributions of mRNA estimations per single cell in the *Travaglini*[3] dataset based on different approaches. Lower triangular matrix shows heat-scatterplots for each pair of approaches, upper triangular matrix shows Spearman's correlation coefficient of these comparisons. Plots in the diagonal show the density for each approach.

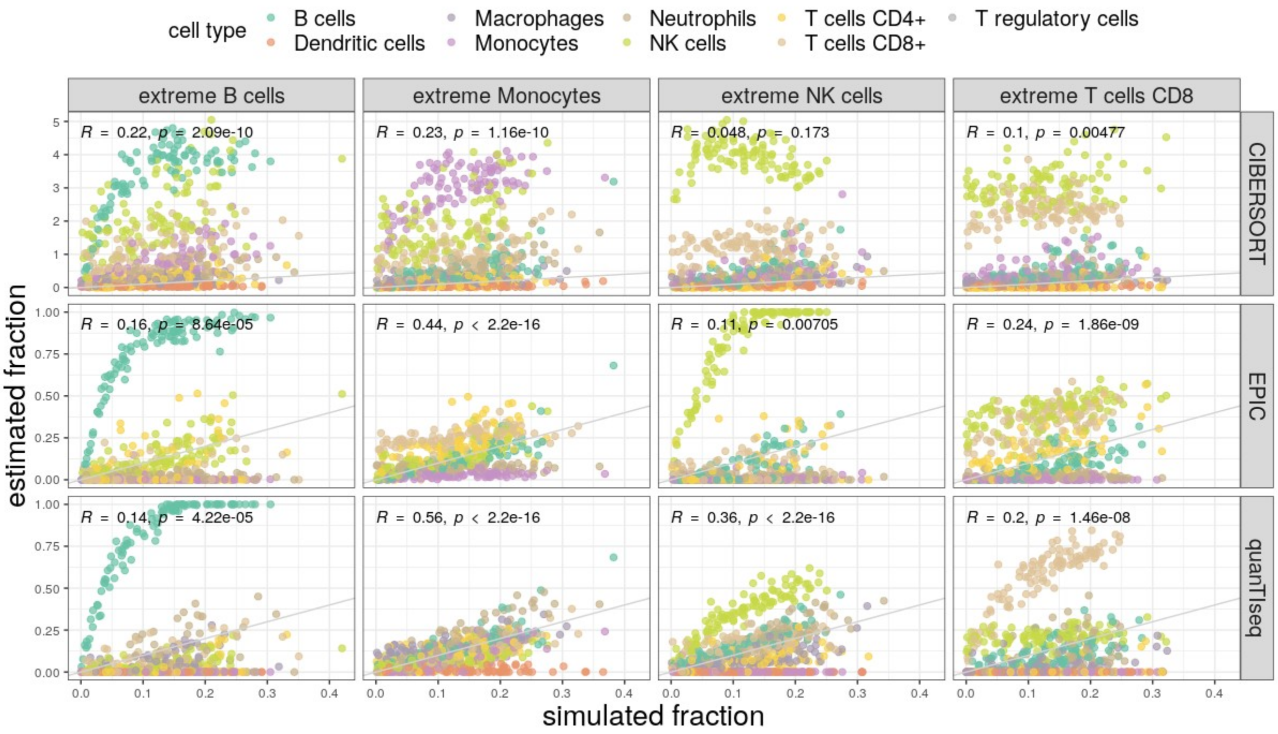


**Supplementary Figure 6:** same setup as in figure 4a of main text, but with added deconvolution results of *EPIC* and *CIBERSORT*.

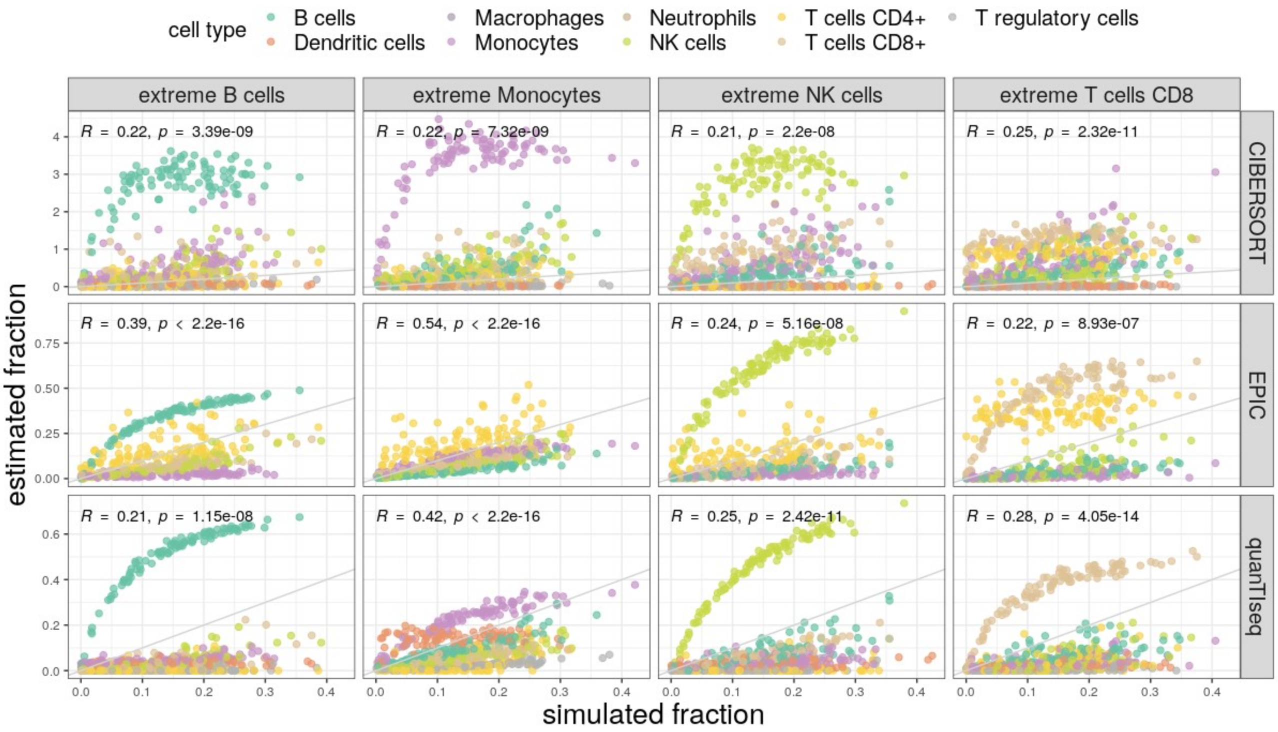


**Supplementary Figure 7:** same setup as in figure 4a of main text, but for *Hao*[1] dataset.

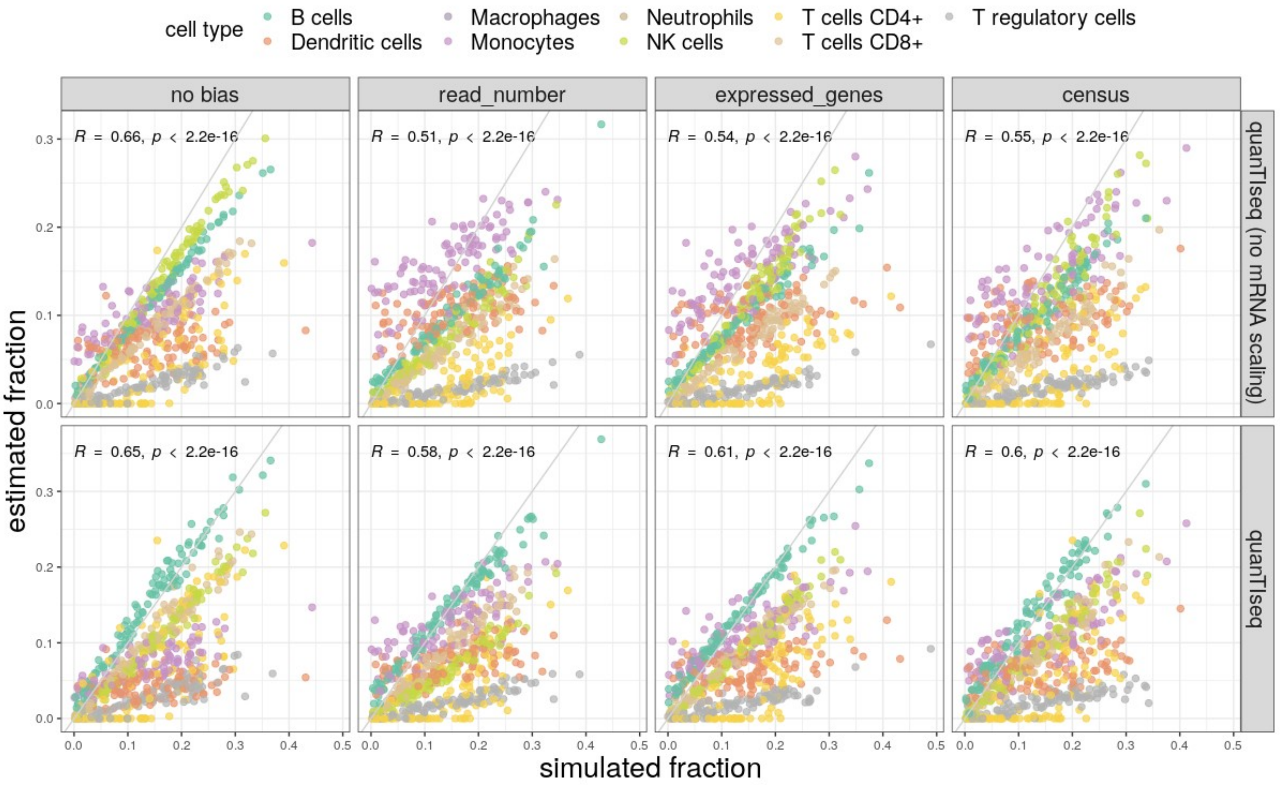


**Supplementary Figure 8:** same setup as in figure 4b of main text, but for *Hao*[1] dataset.

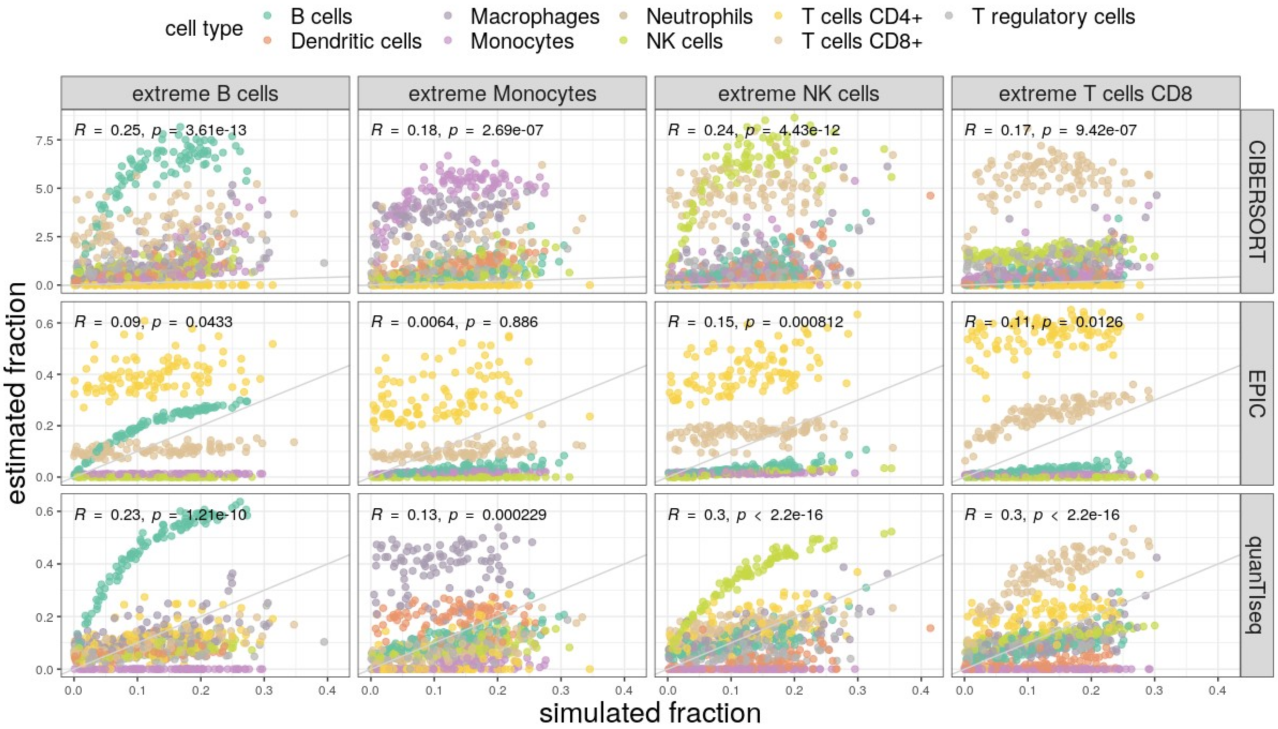


**Supplementary Figure 9:** same setup as in figure 4a of main text, but for *Maynard*[2] dataset.

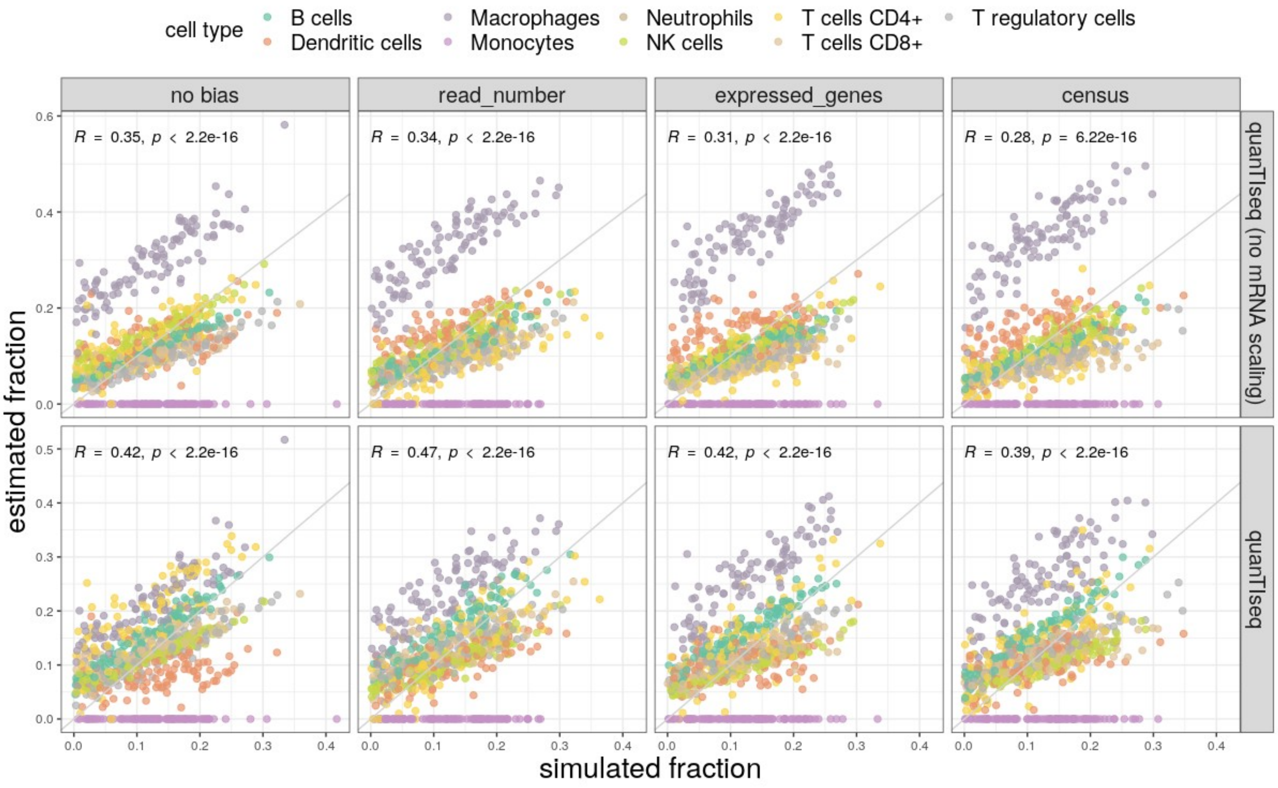


**Supplementary Figure 10:** same setup as in figure 4b of main text, but for *Maynard*[2] dataset.

### Supplementary tables

**Supplementary Table 1:** number of cells per annotated cell type for the three scRNA-seq studies used in this manuscript.

*Table is supplied as separate Excel Sheet.*

**Supplementary Table 2:** names of cell types used for simulations based on three different scRNA-seq studies.

| ***Cell type*** | *Hao*[1] | *Maynard*[2] | *Travaglini*[3] |
|---|---|---|---|
| **B cells** | X | X | X |
| **Dendritic cells** | X | X | X |
| **Macrophages** | | X | X |
| **Monocytes** | X | X | X |
| **Neutrophils** | | | X |
| **NK cells** | X | X | X |
| **T cells CD4** | X | X | X |
| **T cells CD8** | X | X | X |
| **T regulatory cells** | X | X | |

**Supplementary Table 3:** names of cell types used for the simulations based on the Tabula Muris[4] Smart-seq2 and 10x Chromium datasets, using cell type fractions of two true bulk datasets (*Chen*[5,6] and *Petitprez*[7]).

| ***Cell type*** | *Chen (Smart-seq2 based)* | *Chen (10x based)* | *Petitprez (Smart-seq2 based)* | *Petitprez (10x based)* |
|---|---|---|---|---|
| **B cells** | X | X | X | X |
| **T cells CD4** | X | X | X | X |
| **T cells CD8** | X | X | X | X |
| **NK cells** | | | | X |
| **Macrophages** | | | | X |
